# A subpopulation of peripheral sensory neurons expressing the Mas-related G Protein-Coupled Receptor d (Mrgprd) generates pain hypersensitivity in painful diabetic neuropathy

**DOI:** 10.1101/2022.10.27.514066

**Authors:** Dale S George, Nirupa D Jayaraj, Dongjun Ren, Rachel E. Miller, Anne-Marie Malfait, Richard J Miller, Daniela M Menichella

**Author notes:** **CORRESPONDING AUTHOR:** Daniela Maria Menichella, Daniela Maria Menichella, MD., PhD., Department of Neurology and Department of Pharmacology, Feinberg School of Medicine, Northwestern University, Robert Lurie Medical Research Center, Lurie 8-123, 303 E. Superior St, Chicago, IL 60611, USA.

## Abstract

Painful diabetic neuropathy (PDN) is one of the most common and intractable complications of diabetes. PDN is characterized by neuropathic pain accompanied by dorsal root ganglion (DRG) nociceptor hyperexcitability, axonal degeneration, and loss of cutaneous innervation. However, the complete molecular profile underlying the hyper-excitable cellular phenotype of DRG nociceptors in PDN has not been elucidated. This gap in our knowledge is a critical barrier to developing effective, mechanism-based, and disease-modifying therapeutic approaches which are urgently needed to relieve the symptoms of PDN. Using single-cell RNA sequencing we demonstrated an increased expression of the Mas-related G Protein-Coupled Receptor d (Mrgprd) in a subpopulation of DRG neurons in the well-established High-Fat Diet (HFD) mouse model of PDN. *In vivo* calcium imaging allowed us to demonstrate that activation of Mrgprd receptors expressed by cutaneous afferents produced DRG neuron hyper-excitability and oscillatory calcium waves. Furthermore, Mrgprd-positive cutaneous afferents persist in diabetic mice skin. Importantly, limiting Mrgprd signaling or Mrgprd-positive DRG neuron excitability, reversed mechanical allodynia in the HFD mouse model of PDN. Taken together, our data highlights a key role of Mrgprd-mediated DRG neuron excitability in the generation and maintenance of neuropathic pain in a mouse model of PDN. Hence, we propose Mrgprd as a promising accessible target for developing effective therapeutics currently unavailable for treating neuropathic pain in PDN. Furthermore, understanding which DRG neurons cell type is mediating mechanical allodynia in PDN is of fundamental importance to our basic understanding of somatosensation and may provide an important way forward for identifying cell-type-specific therapeutics to optimize neuropathic pain treatment and nerve regeneration in PDN.

## INTRODUCTION

Painful diabetic neuropathy (PDN) is an intractable complication affecting some 25% of diabetic patients^1,2^. It is widely appreciated that the symptoms of PDN include neuropathic pain and small-fiber degeneration^3-6^ involving the degeneration of nerve terminals of the axons of the nociceptive dorsal root ganglion (DRG) neurons that innervate the skin^7,8^. it is well established that peripheral nerve pathology in diabetic patients is characterized by progressive nerve fiber loss with small fibers preferentially affected in early stages with loss of intra-epidermal nerve fibers that can be detected as an early sign in the skin of diabetic patients^9,10^. Neuropathic pain associated with PDN is a debilitating affliction having a substantial impact on patients’ quality of life and health care costs^11^. Despite this prevalence and impact, current therapies for PDN are only partially effective. For example, opioids are not particularly effective in treating neuropathic pain and, given the chronic nature of this syndrome, their use is problematic^12^. Other drugs, such as gabapentinoids and antidepressants produce limited relief in some patients, but the presence of many side effects and their lack of effectiveness in many individuals^13,14^ means that better mechanism-based therapeutic approaches are urgently needed. One critical barrier to developing novel and effective therapy for PDN is that the molecular mechanisms leading to neuropathic pain and to small-fiber degeneration are mostly unknown.

Neuropathic pain is associated with the hyperexcitability of neurons in pain pathways in the absence of appropriate stimuli^4,15,16^. The cells responsible for this phenomenon include DRG nociceptors^4,15,16^. Diabetic patients^17^ and animal models of PDN^18,19^ exhibit sensory neuron hyperexcitability, including spontaneous activity of DRG nociceptor axons^18-20^. Consistent with these findings, our laboratory has shown that reducing the hyperexcitability of DRG nociceptors, identified by the sodium channel Na_v_1.8, which is expressed by 90% of nociceptors^21^, not only reversed mechanical allodynia in the well-established high-fat diet (HFD) mouse model of PDN^6^ but also reversed small-fiber degeneration^22^. It is known that in states of neuropathic pain, DRG nociceptors become hypersensitive to a variety of signaling molecules^9-31,23-28^. Indeed, we demonstrated that activation of the chemokine receptor CXCR4, a G-protein-coupled, seven-span transmembrane receptor (GPCR), was able to increase the excitability of DRG nociceptors in the HFD mouse model of PDN^22^. However, the complete molecular profile underlying the hyper-excitable cellular phenotype of DRG nociceptors in PDN has not been elucidated. The identification of targets able to specifically modulate DRG nociceptor hyper-excitability could help in the further development of novel therapeutic approaches for PDN that could not only treat pain but also reverse small-fiber pathology.

Since nociceptors are a heterogeneous group of neurons^29-35^, accurate and complete classification of the molecular properties of these neurons is imperative if we wish to understand which subtypes are involved in different pathological conditions such as PDN. Recently published single-cell transcriptomic studies have indicated the feasibility of investigating DRG neuron gene regulation at high resolution, with the idea that functional cell type identity will be reflected in the gene expression profile of individual cells within the DRG. Several groups have performed scRNA-seq of rodent and human sensory neurons^29-32,36,37^, facilitating extensive molecular characterization, allowing clustering of DRG neurons and associated non-neuronal cells into distinct subtypes^29-32,36,37^ and clarifying their developmental lineages^37^. Hence, our understanding of the subtypes of DRG sensory neurons is now rapidly evolving through single-cell transcriptomic analysis^29-32,36,37^. However single-cell RNA sequencing and the complete gene expression profile of molecularly distinct DRG cell types, along with the specific genes differentially expressed in each type, have not been addressed in PDN.

We have previously demonstrated that the development of mechanical allodynia was inhibited following the reduction of hyperexcitability in Na_v_1.8-positive DRG neurons in the HFD mouse model of PDN^22^. The subtypes of DRG neurons traditionally linked to mechanical allodynia are C-fibers^38-41^, low-threshold C-mechanoreceptors, and Ad-mechanoreceptors^34,42-45^. However, mechanical allodynia is also mediated by low-threshold Aβ-mechanoreceptors ^34,42^. Traditionally, cutaneous C fibers have been categorized into peptidergic afferents that express substance P or calcitonin gene-related peptide (CGRP), as well as the so-called nonpeptidergic afferents that express Mas-related G Protein-Coupled Receptor d (Mrgprd)^46^. The role of Mrgprd-expressing neurons in mechanical nociception is well established in mice^47,48^. Given that all these neuronal populations express Na_v_1.8 to some degree^21^, our previous studies do not completely deconvolute the nature of the subtypes of neurons within the Na_v_1.8 population that are specifically associated with the occurrence of mechanical allodynia in PDN. Understanding which cell type is mediating mechanical allodynia in PDN is of fundamental importance to our basic understanding of somatosensation and may provide an important way forward for identifying therapeutic targets in this disease.

With this aim in mind, we performed single-cell RNA sequencing of lumbar DRG, so as to capture, in an unbiased fashion, the complete molecular heterogeneity of DRG neurons, and characterized changes in the different DRG cell types. In the well-established High-Fat Diet mouse model of PDN we demonstrated an increased expression of Mrgprd in a subpopulation of DRG neurons in diabetic mice. Mrgprd is an interesting target because is a highly druggable, excitatory G protein-coupled receptor known to influence DRG neuron excitability to mechanical stimuli, expressed solely by the nociceptive neuronal population that extends out into the outermost layer of the skin^48^. Indeed using *in vivo* calcium imaging, we demonstrated that activation of Mrgprd expressed by cutaneous afferents produced DRG neuron hyper-excitability and oscillatory calcium waves. Interestingly Mrgprd-positive cutaneous afferents persist in diabetic mice skin. Furthermore, we were able to reverse mechanical allodynia by genetically limiting Mrgprd signaling or reducing the excitability of Mrgprd expressing DRG neurons. Taken together, our data highlights a key role of Mrgprd-mediated DRG neuron excitability in the generation and maintenance of mechanical allodynia in a mouse model of PDN.

## RESULTS

### Single-cell transcriptomic profiling of dorsal root ganglion (DRG) neurons in the high-fat-diet (HFD) mouse model of painful diabetic neuropathy

In order to understand the mechanisms leading to the DRG nociceptor excitability which underlies neuropathic pain and small-fiber degeneration in PDN, we used the well-established High-Fat-Diet (HFD) mouse model of PDN where mice are fed either a regular diet (RD) or a diet with a high content of fat (HFD) for about ten weeks, during which time they develop mechanical allodynia and small-fiber degeneration^22,49^. Our intention was to discover molecular markers responsible for the hyper-excitable cellular phenotype of DRG neurons as potential novel therapeutic targets. Hence, we identified the complete molecular profile of DRG neurons in their hyper-excitable state using an unbiased transcriptomic comparison of gene expression by lumbar DRG neurons from mice fed an HFD compared mice fed RD. DRG neurons are molecularly defined as distinct subtypes based on the expression of a set of molecular markers^29-32^. To capture the full heterogeneity of DRG neurons and characterize changes in cell types and cell states, we performed single-cell RNA sequencing to understand the complete gene expression profile of molecularly distinct cell types together with the specific genes differentially expressed in each type in the HFD mouse model of PDN.

We prepared viable single-cell suspensions from lumbar DRGs in mice fed a RD or HFD and employed the 10x platform for the single cell capture and barcoding (Fig 1A). We performed comparative clustering (n = 5 mice per group x two rounds; RD 6888 and 6693 cells; HFD 8567 and 6429 cells) and were able to clearly identify different cell types as well as conserved markers in the two conditions investigated. As expected, and in support of the literature on this topic^29-32,36,37^, we were able to separate out neuronal and non-neuronal clusters. Within the neuronal clusters, we were able to identify distinct neuronal subtypes comparable to those described in the literature^29-32,36,37^ (Fig 1B and C). In our comparative analysis of the single-cell data, we were particularly interested in the differential expression of GPCRs as they might represent important drug targets. Mrgprd is an interesting potential drug target because it is a G-protein coupled receptor known to influence DRG neuron excitability to mechanical stimuli^47^. Additionally, Mrgprd is expressed by the nociceptive neuronal population that extends out into the outermost layer of the skin^48^, making it a very accessible therapeutic target for PDN. We confirmed that in RD DRGs, Mrgprd is expressed in a subpopulation of neurons classified as non-peptidergic type1 neurons (NP1)^31,36^ (Fig 1D).

**Figure 1:**
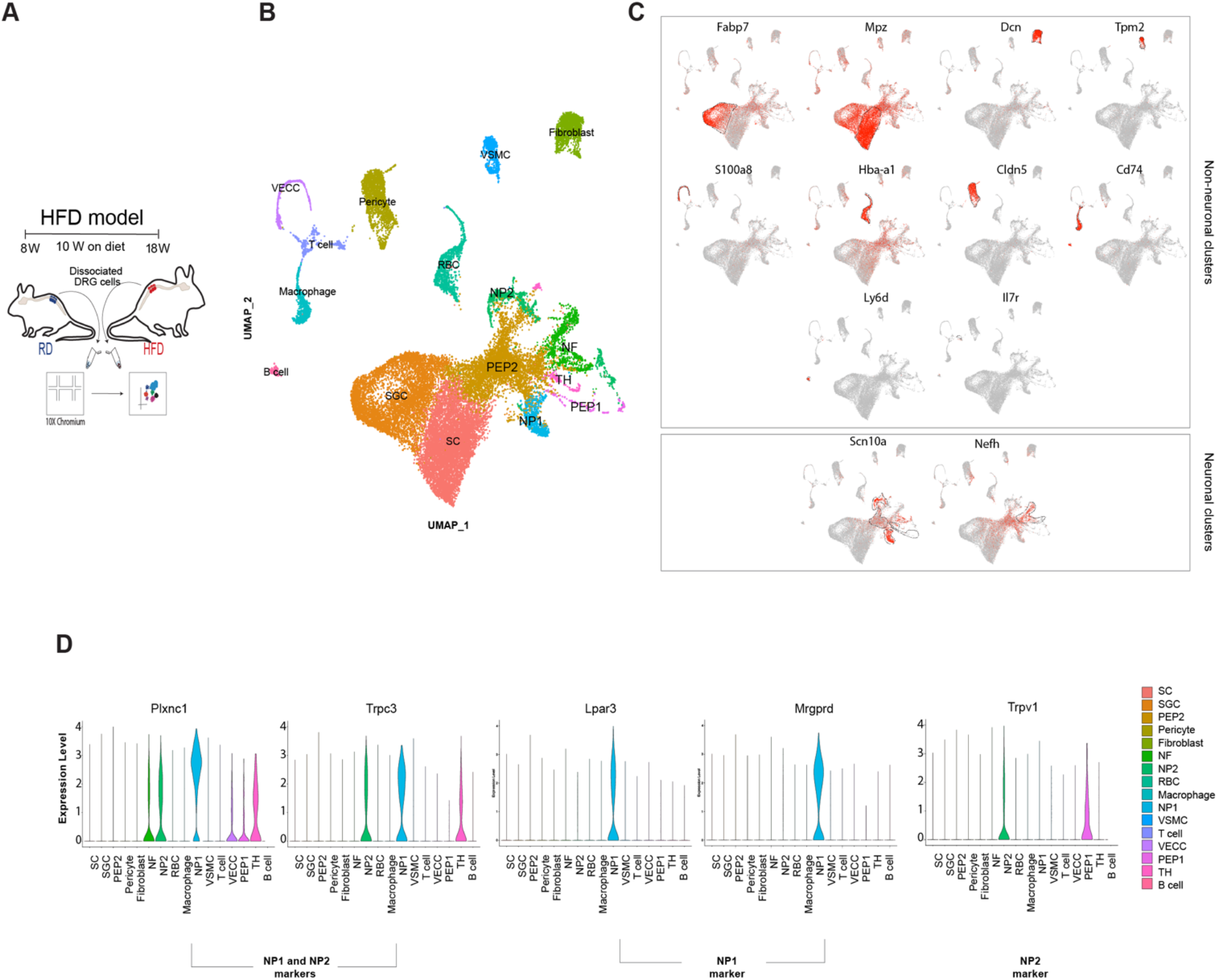
Single-Cell RNA-seq of lumbar DRG neurons in mice fed a RD or a HFD. **(A)** Experimental model describing the two diet groups, isolation of DRG and single-cell capture and barcoding using the 10X Genomics platform. **(B)** Integrated UMAP plot visualizing neuronal and non-neuronal clusters. **(C)** Feature plot indicating the expression of well-known non-neuronal markers and visualization of two broad neuronal markers – Na_v_1.8 and Nefh. **(D)** Violin plot showing expression of well-established markers used to identify the NP1 and NP2 subpopulation. (n = 5 mice per group x two rounds; RD 6888 and 6693 cells; HFD 8567 and 6429 cells).

### Over-expression of Mrgprd receptors in a subpopulation of dorsal root ganglion (DRG) neurons in diabetic mice

Aiming to further evaluate Mrgprd expression when comparing HFD and RD lumbar DRG we performed a comparative analysis of the identified clusters (Fig 2A). The relative proportion of cells in the neuronal clusters indicated an expansion in the peptidergic 1 (PEP1) and non-peptidergic type2 (NP2) clusters (Fig 2B) in the HFD condition. When we compared RD and HFD DRGs, we observed that while there were no differences in the expression of Mrgprd in the NP1 neuronal cluster, Mrgprd was now significantly expressed in NP2 cluster (Fig 2C). The uniquely increased expression of Mrgprd in the NP2 subpopulation could be highly significant in mediating hyperexcitability in these DRG neurons because Mrgprd is an excitatory receptor with considerable constitutive activity^50^. The NP2 population expresses Trpv1, the receptor for capsaicin (Fig 1D and 2D) and we see that there are no differences in the expression of Trpv1 (Fig 2D) in the NP2 population or in the PEP1 population when comparing RD and HFD conditions.

**Figure 2:**
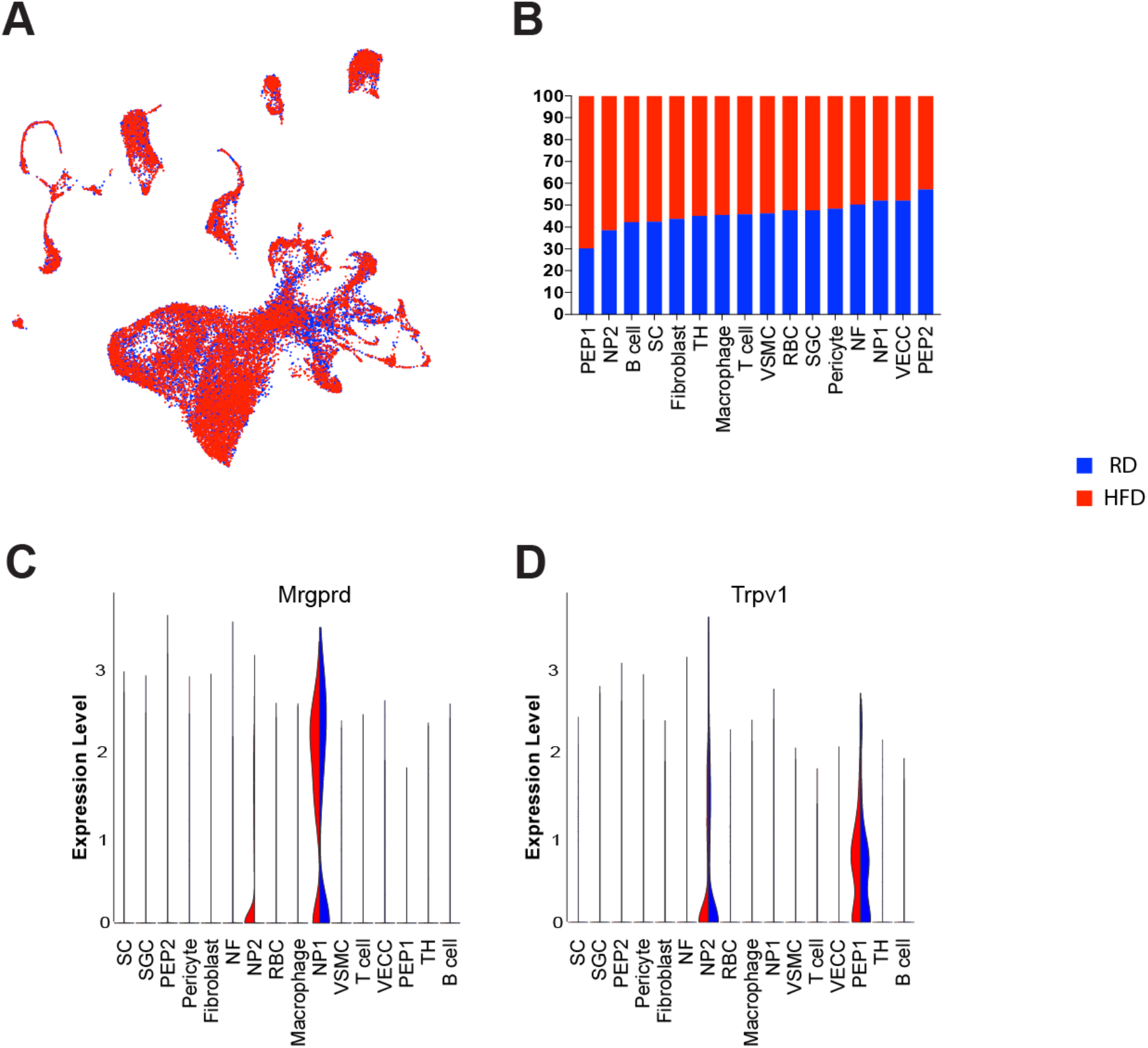
Single-Cell RNA-seq of lumbar DRG neurons reveals over-expression of Mrgprd in mice fed a HFD. **(A)** UMAP grouped by treatment **(B)** Percentage of cells within each neuronal cluster. **(C)** Violin plot indicating changes in gene expression of Mrgprd in the HFD condition. **(D)** Violin plot showing Trpv1 expression in the two diet conditions. (n = 5 mice per group x two rounds; RD 6888 and 6693 cells; HFD 8567 and 6429 cells)

In order to validate the DRG subtypes identified with single-cell RNA seq, we performed RNAscope on frozen sections of lumbar DRG taken from mice fed a RD or a HFD. In particular, we used probes to detect markers characteristic of DRG subgroups. We know that Mrgprd expressing neurons are a subpopulation of the Na_v_1.8 population and do not express the Nefh (Supplemental Fig 1A). Additionally, both in RD and HFD lumbar DRG, we verified the colocalization of Mrgprd and lysophosphatidic acid receptor (Lpar3), a marker of the NP1 DRG subgroups used in previous studies^31^ (Supplemental Fig 1A).

Given the molecular differences in neuronal subpopulations between mouse and human DRG ^51-54^, it was important to validate our potential candidate target using human DRGs. Using RNAscope, we validated the expression of MRGPRD in human DRGs from donor controls and PDN patients provided to us by Anabios. We were able to confirm that MRGPRD is expressed in human DRGs both in control and donors with PDN (Supplemental Fig 1B, 10x magnification). Additionally, as previously demonstrated^51-55^, we confirmed that TRPV1 is more widely expressed in human DRGs as compared to mouse DRGs^53^ both in controls and donors with PDN. Indeed, we observed that TRPV1 was expressed in most nociceptors and co-expressed with MRGPRD (Supplemental Fig 1B, the inset shows a magnified image of a single neuron with the white dashed line demarcating the cell body of a single neuron).

### Activation of Mrgprd at the level of cutaneous afferents produces DRG nociceptor hyperexcitability in diabetic mice

Our single-cell analysis indicated that in diabetic mice Mrgprd is now significantly expressed in the subpopulation of NP2 neurons. Mrgprd is an excitatory receptor, therefore this increased Mrgprd expression could be highly significant in mediating hyperexcitability in these DRG neurons in the HFD model. In order to test this hypothesis, we used *in vivo* calcium imaging as a read-out of neuronal excitability. Given the cellular diversity and functional heterogeneity of DRG neurons^29-32^, we selectively monitored [Ca^2+^]_i_ *in vivo* in Na_v_1.8-positive DRG nociceptors by expressing the [Ca^2+^]_i_ indicator protein GCaMP6^56^ in these neurons (Na_v_1.8-Cre mice^57^ crossed with conditional reporter GCaMP6 mice; Ai96^*flox/flox*^;RCL-GCaMP6s^56^). Mrgprd is expressed in a subpopulation of Na_v_1.8-positive DRG neurons^29-32,36,37^ and, at the transcriptional level, we found no detectable differences in the expression of Na_v_1.8 within either of the two Mrgprd expressing clusters. Na_v_1.8-Cre;Ai96^*flox/flox*^;RCL-GCaMP6s mice were then fed either a RD or HFD for 10 weeks. Laminectomy was performed on anesthetized mice to expose the fourth lumbar (L4) DRG, which contains sensory neurons that innervate the paw^58^.

The *in vivo* physiological setup allowed us to examine DRG neuronal calcium signaling in real-time in response to paw manipulation with mechanical stimuli or application of drugs. To functionally determine changes in Mrgprd expression, we used β-alanine, a known Mrgprd agonist^59^, injected into the paw of mice fed either RD or HFD while monitoring calcium transients *in vivo* in the DRGs (RD: n = 7 mice; HFD: n = 9 mice; average number of neurons within the DRG imaged per animal RD = 197.5 ± 48.67; HFD 242.5 ± 69.16). The number and percentage of responding neurons along with the intensity of these responses were assessed with both vehicle (saline injections) and with β-alanine; representative images are shown in Figure 3A (magnified inset of the whole imaged DRG). While neurons from both RD and HFD mice responded to β-alanine injections into the paw, we found an increase in the percentage of neurons that responded to the stimulus in HFD mice (Fig 3B, RD 2.30 ± 0.51; HFD 5.81 ± 2.9, *p* = 0.008, white circles indicate neurons, and the yellow circles identify neurons that responded to the stimulus at a given time).

**Figure 3:**
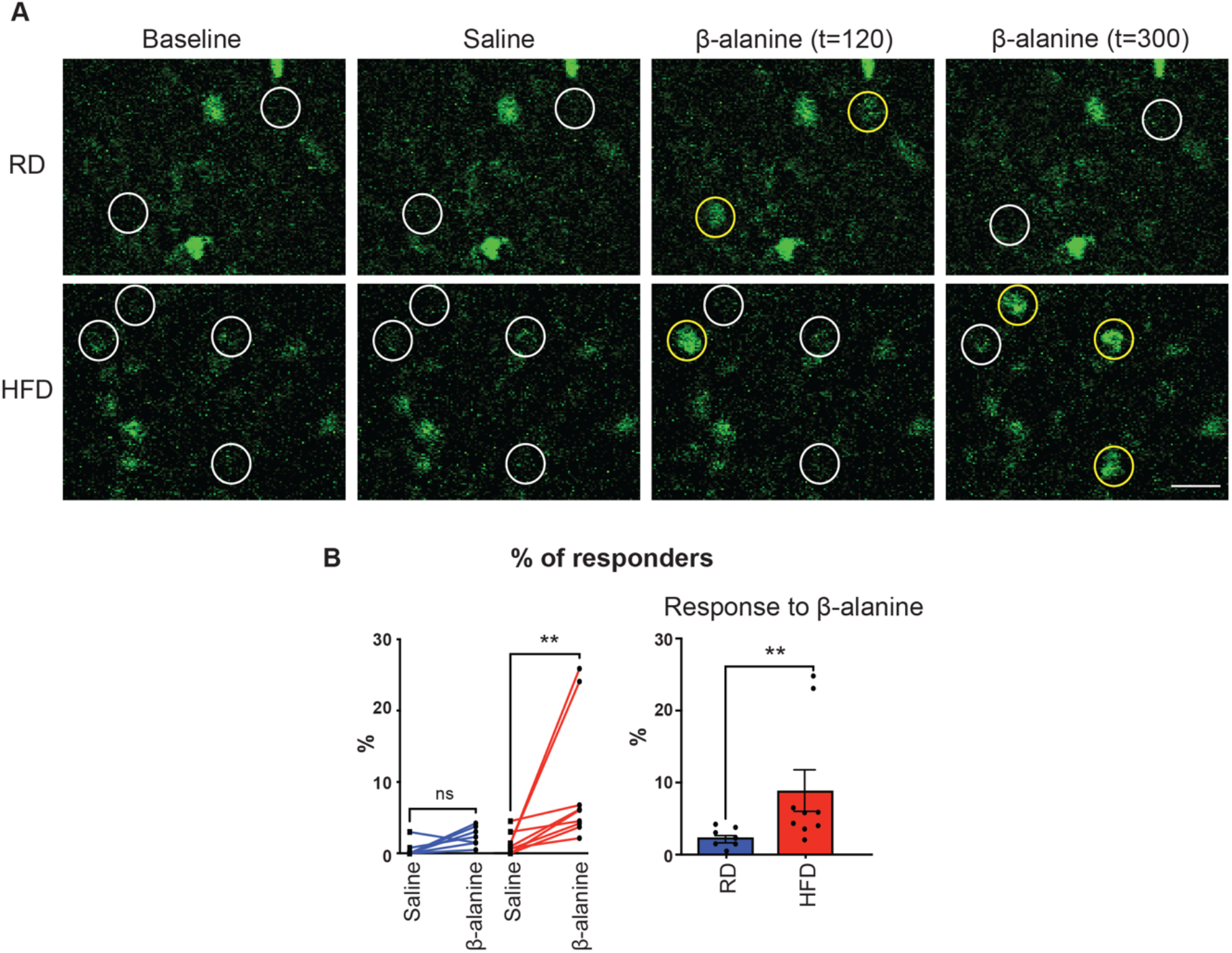
Increased percentage of in vivo calcium responses to β-alanine in Na_v_1.8-expressing DRG neurons from mice fed a HFD. **(A)** Representative magnified images of whole DRG taken from RD (top) and HFD (bottom) mice showing neurons identified by white circles at baseline and during administration of saline. A neuron that responds to the stimulus is identified by a yellow circle at t = 120 seconds and at t = 300 seconds. **(B)** Paired analysis showing the percentage of responses to saline (square) and β-alanine (circle) treatment in RD (blue) and HFD (red) and percentage of neurons responding to β-alanine, each black circle represents one animal. RD (n = 7 mice); HFD (n = 9 mice) (average number of neurons within the DRG imaged per animal RD = 197.5 ± 48.67; HFD 242.5 ± 69.16). \*\**p* < 0.01.

### Activation of Mrgprd at the level of cutaneous afferents in the skin produces oscillatory calcium waves in a subpopulation of Mrgprd expressing DRG neurons in diabetic mice

In the HFD condition, our single-cell RNA sequencing revealed two clusters that expressed Mrgprd, NP1, and NP2 neuronal clusters (Fig 2C). We wished to determine the significance of these observations in producing DRG neuron hyperexcitability. Interestingly, we were able to distinguish two distinct patterns of calcium signaling as a result of activating Mrgprd expressed by DRG neurons. Among the DRG neurons taken from the HFD group, many responses were similar in amplitude to those seen in RD (Fig 4A, E; RD 1.56 ± 0.14, HFD 1.78 ± 0.08, *p* = 0.19). RD (n = 7 mice); HFD (n = 9 mice) (average number of neurons within the DRG imaged per animal RD = 197.5 ± 48.67; HFD 242.5 ± 69.16). However, there was also a Mrgprd expressing subpopulation of neurons from the HFD group in which activation of Mrgprd receptors with β-alanine injection in the paw produced very marked long-lasting oscillatory calcium responses, something that was rarely observed in the RD group. (Fig 4B). Neurons in the RD group start to respond within 20 seconds after administration of β-alanine (t = 0 - 99 seconds of baseline reading, β-alanine injected at t = 100 seconds) and most of the calcium peaks are no longer detected beyond 300 seconds (Fig 4A, B; RD 10.67%). Similarly, neurons in the HFD also start to respond 20 seconds after the administration of β-alanine but, interestingly, when compared with the RD group, about double the percentage of neurons in the HFD continued to respond even after 300 seconds (Fig 4A, B; HFD 23.17%). Figure 4C shows all the responses from an individual animal as an illustration of the oscillatory calcium responses detected in the HFD group (Fig 4C, grey trace indicates the average response).

**Figure 4:**
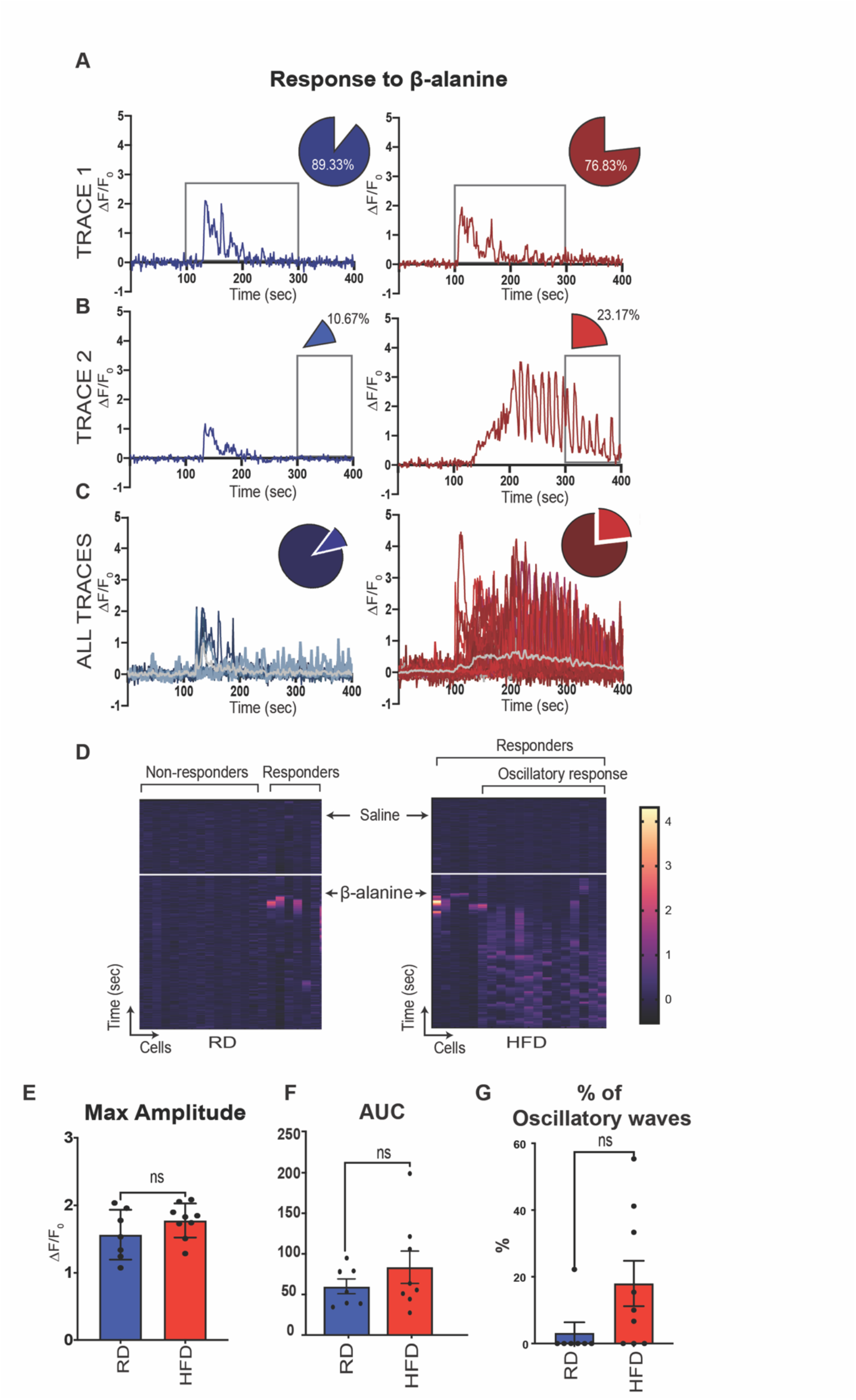
Oscillatory in vivo calcium responses in mice fed a HFD. **(A)** Trace from a single neuron in RD (blue) and HFD (red) indicating change in fluorescence intensity (ΔF/F_0_) over time in seconds (sec). **(B)** Trace from a single neuron from RD and HFD in response to β-alanine injection. **(A-B)** The box shows the duration of response to β-alanine in both RD and HFD. The percentage of neurons responding within this duration is indicated by the pie chart. **(C)** All responses from representative RD and HFD mice. The gray line indicates the average change in fluorescence intensity over time. **(D)** Heat map of 20 neurons shown as a function of time and their response to saline and β-alanine injections (arrows indicate the time of injection) **(E)** Graphs showing maximum amplitude, area under the curve (AUC) and the % of oscillatory waves. RD (n = 7 mice); HFD (n = 9 mice) (average number of neurons within the DRG imaged per animal RD = 197.5 ± 48.67; HFD 242.5 ± 69.16).

The heat map visualization enables us to appreciate this oscillatory behavior of neurons in the HFD (Fig 4D, 20 neurons from a representative animal fed an RD or HFD). So, while a population of β-alanine responsive neurons in the HFD responded with calcium transients similar in amplitude and duration to the neurons in the RD (Fig 4E, F), a percentage of β-alanine responsive neurons had a different profile, with oscillatory waves (Fig 4D, G, RD = 3.17 ± 3.17 HFD = 17.99 ± 6.81, *p* = 0.07) and longer duration of response. Focusing on the Na_v_1.8-positive DRG neurons responding to β-alanine, we further determined their response to capsaicin (Supplemental Fig 2) and found no statistically significant differences when comparing RD and HFD groups. Taken together these results support the idea that there are indeed two different subpopulations of Mrgprd-expressing neurons, and one subtype might be more important in mediating the hyperexcitability observed in the HFD model of PDN.

### Mrgprd-positive cutaneous afferents persist in diabetic mice skin

Previously our lab has shown that there is a loss of the Na_v_1.8-positive intraepidermal nerve fibers in the HFD condition^22^. Since in our *in vivo* calcium imaging experiments involved injecting β-alanine into the paw, it seemed counterintuitive that we were able to detect responses within the DRG if these neurons failed to innervate the skin. Hence, we determined the status of the Mrgprd fibers in the skin. For this, we used a Mrgprd-eGFP reporter mouse (MrgprdΔ EGFPf)^48^ and we induced PDN by feeding either a RD (n = 4 mice) or a HFD (n = 3 mice) for ten weeks. The mice gained weight (Supplemental Fig 3A) and became glucose intolerant ten weeks after the onset of HFD (Supplemental Fig 3B, and 3C). We next examined skin samples of Mrgprd-eGFP mice using confocal microscopy. We observed that Mrgprd-positive cutaneous afferents normally crossed the intra-epidermal junction and innervated the outer layer of skin even in the HFD condition, with no statistically significant difference in the intra-epidermal nerve fiber (IENF) density (expressed as the number of nerves crossing the epidermal-dermal junction as a function of length) between RD (0.08 ± 0.01) and HFD (0.07 ± 0.02) mice (Fig 5A, B). Thus, it is possible that these remaining or regenerating cutaneous afferents expressing the excitatory Mrgprd receptors could be responsible for mediating mechanical allodynia in this model. Given that these Mrgprd receptors remain in the outer layers of the skin, they become a remarkably interesting and accessible target for an antagonist that could be applied topically to the skin.

**Figure 5:**
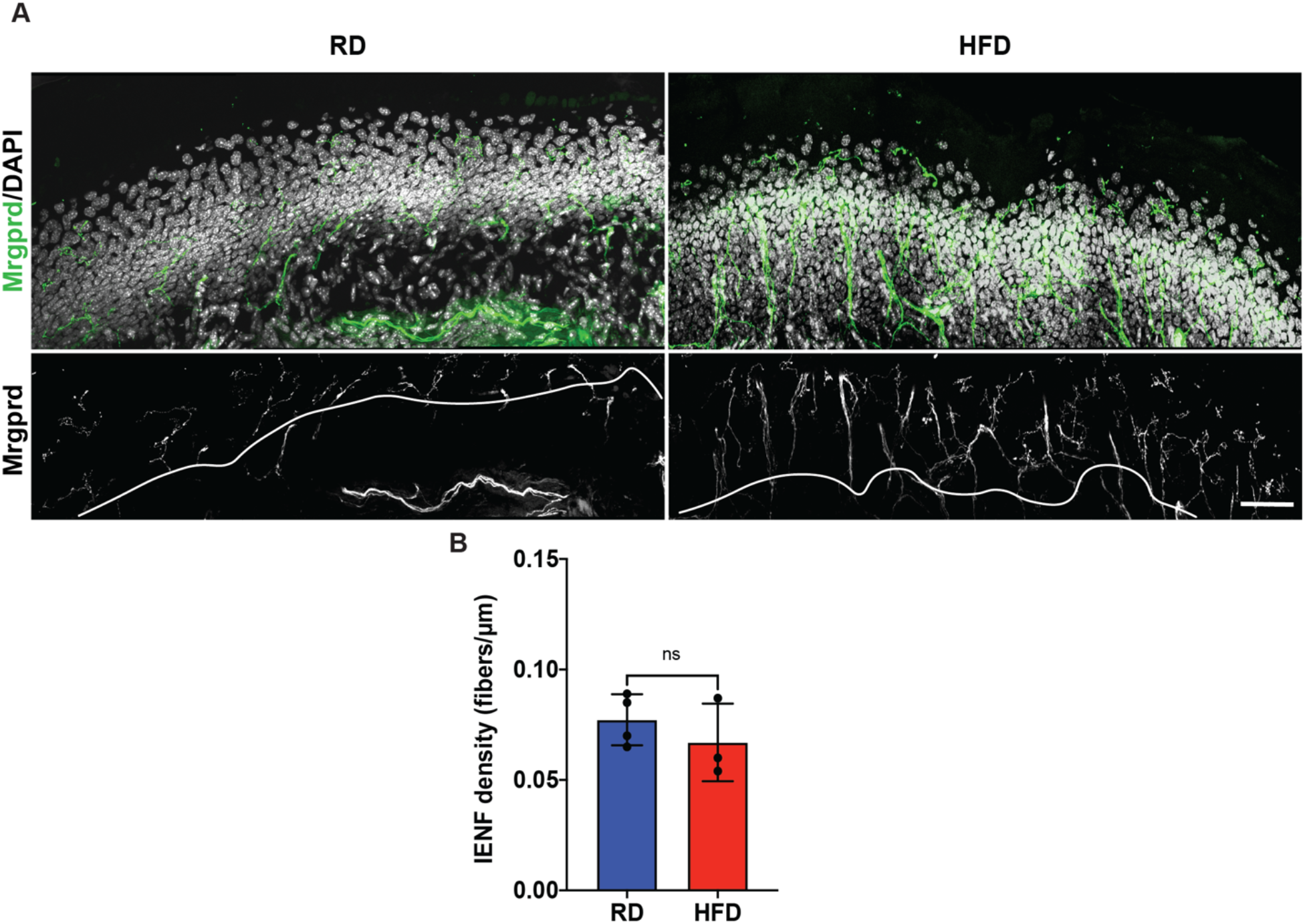
Persistence of the Mrgprd afferents in the hind-paw of animals fed a HFD. **(A)** Confocal micrographs of the glabrous skin of mice fed either a RD or HFD showing Mrgprd expressing neuronal afferents labeled with GFP (green) and nuclear marker DAPI (gray). The bottom panel indicate the intra-epidermal region with the white line demarcating the intra-epidermal junction and the Mrgprd expressing fibers are shown in gray. Scale bar represents 50μm. **(B)** Quantification of the Mrgprd intraepidermal nerve fiber density in RD and HFD (RD n=4 mice, HFD n=3 mice; 5 sections for each group).

### Mrgprd receptors are necessary for the establishment of static mechanical allodynia in diabetic mice

To make a compelling case for MrgprD receptors as a drug target for PDN, we must demonstrate that manipulating these receptors reduces PDN symptoms. Thus, we examined the effects of reducing the expression of Mrgprd receptors in the development of pain hypersensitivity in the HFD mouse model of PDN using mice in which the Mrgprd receptor had been deleted and replaced with a gene for a fluorescent protein, EGFPf (MrgprdΔ EGFPf)^48^. First, we induced PDN in MrgprdΔ EGFPf mice by feeding them either a RD or HFD. Both MrgprdΔ EGFPf heterozygous and homozygous mice fed HFD developed obesity (Supplemental Fig 3A) and glucose intolerance (Supplemental Fig 3B and 3C) like wild-type mice, demonstrating that Mrgprd receptors deletion does not alter their response to the changed metabolic profile prevailing in the HFD. We then tested both Het and Homo mice for mechanical allodynia (RD-Het n = 10 mice; RD-Homo n = 8 mice; HFD-Het n = 17 mice; HFD-Homo n = 16 mice) using von Frey withdrawal threshold measurements, as described elsewhere^22,49,60^. Deletion of the Mrgprd receptors did not change the withdrawal thresholds or result in mechanical allodynia compared with the heterozygous mice fed a RD (Fig 6A; RD-Het 0.80 ± 0.93 v/s RD-Homo 0.70 ± 0.35, *p* =>0.99) indicating that Mrgprd deletion did not alter mechanical sensation in otherwise metabolically normal mice. We have previously shown that mice fed an HFD developed mechanical allodynia six weeks after commencement of the diet. In Het mice fed the HFD for 10 weeks, the withdrawal threshold was significantly reduced compared to RD-Het mice (RD-Het 0.80 ± 0.93 v/s HFD-Het 0.13 ± 0.24; *p = 0*.*0001*), indicating mechanical allodynia in this group (Fig 6A). We then determined whether deleting the Mrgprd receptor would prevent the development of mechanical allodynia in HFD mice. We saw that there was a significant improvement in the withdrawal threshold between HFD-Het and HFD-Homo mice (Fig 6A; HFD-Het 0.13 ± 0.24 v/s HFD-Homo 0.33 ± 0.57, *p* = 0.03) indicating that deleting the Mrgprd receptors prevented the establishment of mechanical allodynia in this model of PDN.

**Figure 6:**
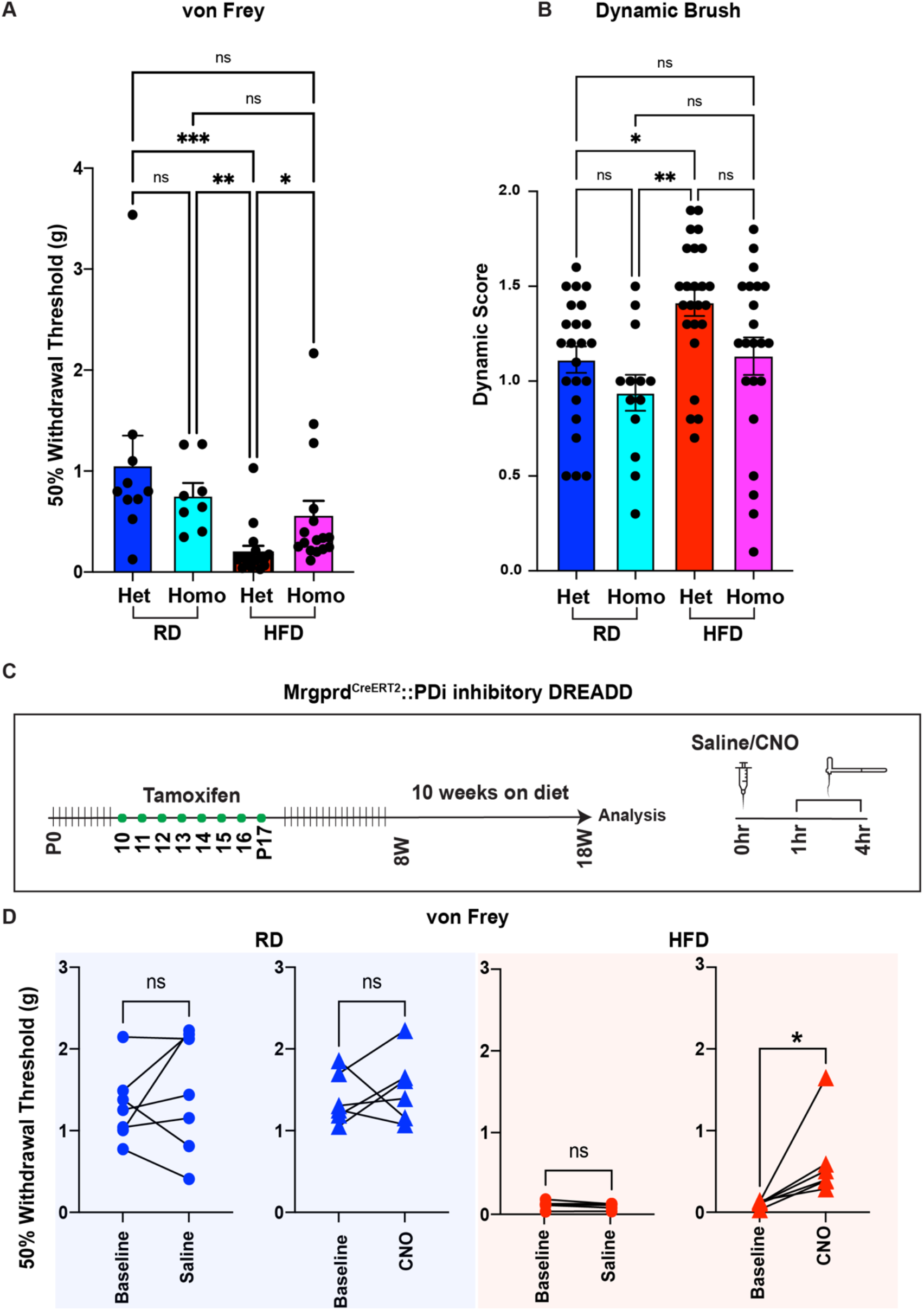
The Mrgprd expressing neurons and the Mrgprd receptor are crucial in mediating mechanical allodynia. **(A)** von Frey testing of heterozygous (+/-) and homozygous (-/-) Mrgprd mice to evaluate static mechanical allodynia (RD-Het n = 10 mice; RD-Homo n = 8 mice; HFD-Het n = 17 mice; HFD-Homo n = 16 mice) **(B)** dynamic brush assay to evaluate the contribution of Mrgprd receptor to dynamic brush allodynia. (RD-Het n = 10 mice; RD-Homo n = 8 mice; HFD-Het n = 17 mice; HFD-Homo n = 16 mice) **(C)** Schematic showing the generation of Mrgprd-CreERT2 crossed with hM4 DREADD mice **(D)** von Frey testing of RD and HFD Mrgprd-CreERT2;hM4D mice at baseline followed by von Frey after acutely administered with either saline (circles) or CNO (triangles). RD saline n=7 mice, RD CN0 n=6 mice, HFD saline n=6 mice, HFD CNO n=6 mice. \*\*\**p* < 0.001, \*\**p* < 0.01, \**p* < 0.05

To further characterize the pain phenotype, we tested for dynamic allodynia using the brush test^61-63^ in both Het and Homo mice fed either RD (RD-Het n = 23 mice; RD-Homo n = 13 mice) or HFD (HFD-Het n = 24 mice; HFD-Homo n = 22 mice) for ten weeks. We observed that mice fed HFD developed dynamic allodynia at ten weeks on diet (Fig 6B; RD-Het 1.20 ± 0.33 v/s HFD-Het 1.45 ± 0.34 *p* = 0.04; RD-Homo 1.00 v/s HFD-Het *p* = 0.002). We observed that homozygous animals fed a HFD did not show an improvement in the dynamic score compared with the HFD-Het mice (HFD-Het v/s HFD-Homo 1.20 ± 0.46 *p* = 0.18) demonstrating that Mrgprd receptors are necessary for the establishment of static but not brush (dynamic) mechanical allodynia in the HFD mouse model of PDN.

### Reducing Mrgprd-positive DRG neuron excitability reversed mechanical allodynia in diabetic mice

Given that Mrgprd receptors appear to be necessary for the establishment of mechanical allodynia, targeting Mrgprd receptors is a promising approach to treating neuropathic pain in PDN. Alternatively, one could target the hyperexcitability of Mrgprd-positive DRG neurons. To test the hypothesis that Mrgprd-positive DRG neuron hyperexcitability has a role in mediating mechanical allodynia in the HFD mouse model of PDN, we used a chemogenetic platform genetically introducing Designer Receptors Exclusively Activated by Designer Drug (DREADD) receptors into Mrgprd-positive DRG neurons. As previously described^22^, we used an inhibitory DREADD receptor based on an engineered muscarinic acetylcholine receptor M_4_ (PDi), which works via activation of the inhibitory G_i/o_ protein pathway ^64^. Activation of this receptor with the small molecule agonist clozapine-N-oxide (CNO) or its metabolite clozapine inhibits neuronal activity (for review ^65-67^). We expressed inhibitory hM_4_ DREADD (PDi) receptors in Mrgprd-positive DRG neurons by crossing Mrgprd-CreERT2 mice^68^ with a mouse line that enables the conditional expression of DREADD receptors (hM4Di, Fig 6C)^66^.

As expected, we observed that HFD Mrgprd-CreERT2;hM4Di mice had significantly lower baseline withdrawal threshold for mechanical stimulation compared to animals on RD (Suppl Fig 4A RD 1.25 ± 0.38 v/s HFD 0.11 ± 0.04, *p* < 0.0001). We then evaluated the consequences of reducing Mrgprd-positive DRG neuron excitability on mechanical allodynia in the HFD model by von Frey analysis one hour after a single intraperitoneal ip injection of either saline or CNO (10 mg/kg) (RD saline n=7 mice, RD CN0 n=6 mice, HFD saline n=6 mice, HFD CNO n=6 mice). We saw no changes in the withdrawal threshold when animals fed a RD were injected with either saline (Fig 6D; RD-baseline 1.30 ± 0.45 v/s RD-saline 1.48 ± 0.73 *p* = 0.58; n = 7 mice) or CNO (RD-baseline 1.39 ± 0.31 v/s RD-CNO 1.52 ± 0.42 *p* = 0.69; n = 6 mice), whereas we saw a significant improvement in the withdrawal threshold when HFD animals were treated with CNO (Fig 6D; HFD-baseline 0.12 ± 0.05 v/s HFD-saline 0.10 ± 0.04 *p* = 0.22; HFD-baseline 0.11 ± 0.04 v/s HFD-CNO 0.64 ± 0.51 *p* = 0.03). The withdrawal threshold subsequently returned to baseline four hours after injection (Supplemental Fig 4B). This data demonstrates that specifically modulating Mrgprd-positive DRG neuron excitability via Gi-coupled GPCRs can reverse mechanical allodynia in the HFD mouse model of PDN.

## DISCUSSION

The results of our experiments demonstrated that the excitatory Mas-related G protein-coupled receptor d (Mrgprd) plays a key role in the generation and maintenance of DRG nociceptor hyperexcitability underlying neuropathic pain in PDN. Using single-cell RNA sequencing we demonstrated increased Mrgprd expression in a subpopulation of non-peptidergic DRG neurons in the HFD mouse model of PDN. These changes in Mrgprd expression were shown to have functional significance in producing DRG neuron excitability underlying PDN. Indeed, activating Mrgprd receptors in the periphery, at the level of the cutaneous afferents, produced DRG neuronal hyper-excitability and associated oscillatory calcium waves in diabetic mice. Importantly, limiting Mrgprd signaling or Mrgprd-positive DRG neuron excitability, reversed mechanical allodynia in the HFD mouse model of PDN. Given the observation that Mrgprd-expressing nerves also remain in the skin during PDN, these studies indicated that Mrgprd-mediated excitability is an accessible target for developing effective therapeutics which are currently unavailable for PDN patients^13^.

Mrgprd is a promising therapeutic target. First, Mrgprd is a G protein-coupled receptor that has often been used as a druggable target^69-71^. Second, Mrgprd is an excitatory receptor with significant constitutive activity^47^ so its overexpression may increase neuronal excitability of DRG neurons even in the absence of an activating ligand. The constitutive activity of the receptor makes it an ideal candidate for generating spontaneous electrical impulses which is a clinical symptom that is commonly observed in PDN^72,73^; hence blocking this activity using an inverse agonist should produce analgesic effects in PDN. Third, Mrgprd is known to influence DRG neuron excitability in response to mechanical stimuli^68,74,75^. Finally, Mrgprd is expressed by the nociceptive neuronal population that extends out into the stratum granulosum, the outermost layer of the skin^47,48^ making it a very accessible therapeutic target for PDN. While Mrgprd, is predominately expressed sensory neurons in dorsal root ganglia (DRG) and trigeminal ganglia (TG) of animals, including humans^61,76^ there are also reports that Mrgprd is expressed outside the nerve system including aortic endothelial cells^77,78^, mouse primary neutrophils^79^ and mouse intestinal tract^80^. If these observations can be confirmed then, from a translational perspective, an ideal therapy for PDN might involve Mrgprd modifying drugs applied topically. Indeed, topical applications for treating PDN are very appealing as they should bypass drug side effects associated with systemic interventions. However, one limitation of our study is that we have used a genetic approach to demonstrate the role of Mrgprd in mediating neuropathic pain in the HFD mouse model of PDN as drugs for doing this are unavailable. The pharmacology of Mrgprd has not been extensively investigated, although an inverse agonist MU-6840^50^ has been reported in a single report in the literature which might be an ideal candidate for examining models of PDN pain.

Numerous single-cell RNA-seq studies of human DRGs have demonstrated that there are several significant molecular differences between rodent and human DRG neuron subtypes^51-55^. Integrative analysis with single-cell RNA-seq data comparing human and mouse DRGs revealed broad conservation of known DRG and/or nociceptor-enriched genes (e.g., *P2XR3* [P2X3 receptor], *SCN10A* [Na_v_1.8], *SCN11A* [Na_v_1.9], *NTRK1* [TrkA], and *MRGPRD* [MRGPRD]) across mouse and human DRGs^52^, although the relative patterns of co-expression of markers by different subsets were different in mice relative to humans^51^. Given the potential molecular differences in neuronal subpopulations between mouse and human DRG^51-54^, it is crucial to validate the molecular mechanisms underlying neuropathic pain discovered in mice using human tissue if they are to be successfully translated to humans^81^. We validated the expression of MRGPRD in human DRGs from donor controls and PDN patients (Supplemental Fig 1B). As previously demonstrated^51-55^, we observed that TRPV1 was expressed in most nociceptors and co-expressed with MRGPRD (Supplemental Fig 1B inset), making the distinction between peptidergic and non-peptidergic DRG neurons previously demonstrated in mice^46^ less clear in human DRGs^51-55^. Mrgprd is part of the large Mas-related G coupled receptors (Mrgprs) superfamily of receptors, Mrgprd being conserved across rodents and humans. However, most recently differences in Mrgpr expression patterns have been demonstrated between rodents and human DRG^82^. The transcriptional heterogeneity among DRG neuronal clusters is an important contributor to the functional specificity and responses to cutaneous stimuli of specific neuronal subtypes in both mice and humans. While the role of Mrgprs both in mediating non-histaminic itch and the excitability of polymodal nonpeptidergic nociceptors to mechanical in rodents has been clearly delineated^47^, the pharmacological characterization and insights into the physiological roles of the majority of Mrgprs in different species, including humans, remains to be completely established. Several peptides and small molecules have been proposed as ligands for Mrgprs^83-85^. However, many of these ligands interact with multiple receptor types^82^. Therefore, future screening for molecule MrgprD antagonist or inverse agonist activity has to be performed using high-throughput electrophysiology or calcium image studies employing human dorsal root ganglion (DRG) neurons or nociceptors derived from human induced pluripotent stem cells (iPSCs) if they are to be successfully translated to humans^81^.

Several groups have performed scRNA-seq of rodent sensory neurons^29-32,36,37^, facilitating their molecular characterization, allowing clustering of DRG neurons and associated non-neuronal cells into distinct subtypes^29-32,36,37^ and delineating their developmental lineages^37^. ScRNA-seq has enabled the potential for studying gene regulation at high resolution, commensurate with the functional cell type identity that will be reflected in the gene expression profile of individual cells within DRG. With this approach, we can now begin to describe gene expression changes within populations that accompany disease states. Although characterization of the gene expression profiles in injured DRG has been extensively studied through bulk RNA sequencing, including transcriptomic analysis with bulk RNA sequencing of DRG from donor control and with painful diabetic neuropathy^86^, these data do not clearly delineate which specific cell types represent these transcriptional changes. Now, scRNA profiling of DRG neurons has been performed in rodents and primates^87^ with chronic pain. For example, Hu et al. performed scRNA-seq on DRG neurons from control mice and mice undergoing sciatic nerve transection^88^. An increase in the transcription of genes associated with cell death and alterations in pathways related to neuropathic pain was observed in the non-peptidergic (NP) neuronal population. This heterogenous injury-induced transcriptional alteration within neuronal subtypes suggests that there are intrinsic differences in the genetic response to injury between subtypes of DRG neurons. To understand the dynamic changes at single-cell resolution during the development of neuropathic pain, scRNA-seq of DRG neurons at different time points after spared nerve injury (SNI) was performed^89^.

In addition to the expected neuronal clusters, three additional clusters were observed after SNI. These clusters had high expression of Atf3, Gal, and other nerve-injury-regulated genes. Interestingly, a cluster expressing Atf3/Mrgprd appeared after 24 hours and transcriptomic changes within this cluster led to changes in neuronal phenotype within two days after injury, highlighting that distinct neuron types respond differently to injury and that injured Mrgprd-expressing neurons have particularly high reprogramming capabilities.

In the present study, we used single-cell RNA sequencing to uncover that Mrgprd is significantly expressed in the non-peptidergic type 2 (NP2) DRG neuron subtype in diabetic DRGs. The role of Mrgprd-expressing neurons in mechanical nociception is well established in mice^47,48^. Using in vivo calcium imagining we were able to distinguish two distinct patterns of calcium signaling as a result of activating Mrgprd expressed by DRG neurons. Indeed we were able to demonstrate that activation of the Mrgprd receptors in the periphery with its known agonist, β-alanine, induced characteristic oscillatory calcium waves only in one subgroup of Mrgprd expressing neurons in diabetic mice, indicating that the two distinct subgroups of non-peptidergic DRG neurons expressing Mrgprd had different physiological proprieties which may be relevant in mediating neuropathic pain in PDN. However, our study does not completely deconvolute the nature of the subtypes of DRG neurons expressing Mrgprd that are specifically associated with the occurrence of mechanical allodynia in our mouse model of PDN. Future studies designed to activate or silence selectively the NP1 or the NP2 subtype of non-peptidergic neurons combined with behavioral assessments are necessary to address these questions.

One of the main hallmarks of PDN is small-fiber degeneration ^3,4^, particularly a “dying back” axonopathy that affects the smallest axons of the peripheral nervous system: the dorsal root ganglion (DRG) nociceptor axons ^7,8^. Previously our lab has shown that there is a loss of the Na_v_1.8-positive intraepidermal nerve fibers in the HFD condition^22^. Here we demonstrated that Mrgprd-positive cutaneous afferents normally cross the intra-epidermal junction and innervate the outer layer of skin even in the HFD condition. It is possible that these cutaneous Mrgprd expressing afferents could be responsible for carrying mechanical allodynia in this model. Indeed, in the HFD model, we observed an increase in the percentage of neurons responding with calcium transients upon activation of Mrgprd receptors expressed in cutaneous nerve terminals by its known agonist β-alanine injected into the skin. Given that these Mrgprd receptors remain in the outer layers of the skin, they represent an interesting and accessible target for an antagonist/inverse agonist that could be applied topically. Indeed, it is known that Mrgprd-expressing fibers specifically innervate the stratum granulosum of the epidermis^48^. One limitation of our study is that we were not able to establish if Mrgprd positive afferents innervating the outermost layer of the skin even in the diabetic condition are nerves that are spared by degeneration or represent newly regenerating fibers. Several studies in rodent models of PDN^90^ and in PDN patients have now demonstrated an increased regeneration of intraepidermal nerve fibers (IENFs)^91,92^. Indeed, increased regeneration of IENFs was able to distinguish between patients with painful and non-painful diabetic neuropathy^92^. Another study reports that increased regenerating IENFs were distinctly associated with ongoing burning pain in patients with diabetes^93^. We are currently exploring the status of Mrgprd positive afferent and their regenerative properties in the skin of rodent models of PDN and in clinically well-characterized patients with PDN, to enhance the translational validity of Mrgprd as a therapeutic target.

Interestingly, using our in vivo two-photon calcium imaging setup that allows visualization of calcium signals in lumbar DRGs in response to manipulation of the mouse paw with either mechanical stimuli or with the application of drugs, in diabetic mice we were able to uncover long-lasting intracellular calcium oscillations in a subset of Mrgprd expressing DRG neurons when activated with their known agonist β-alanine at the periphery. Intracellular calcium (Ca2+) oscillations are an ubiquitous signaling mechanism, occurring in many cell types and controlling a wide array of cellular functions^86,94-97^. Ca2+ oscillations are proposed to convey information in their amplitude and frequency, leading to the activation of specific downstream targets^98-100^. Our study is to our knowledge the first report of long-lasting intracellular calcium oscillations in DRG neurons in the contest of PDN or other painful peripheral neuropathies. Devor and colleagues have shown that axotomy increases the population of DRG neurons exhibiting burst discharges triggered by subthreshold oscillations, and increased ectopic discharges associated with neuropathic pain^101,102^. Oscillatory Ca^2+^ signals generated by periodic release of Ca^2+^ from intracellular stores and/or voltage-dependent Ca^2+^ influx are often utilized to regulate different stages of neural development and are implicated in driving numerous genetic programs including proliferation, and differentiation^103,104^. It is feasible to hypothesize that Mrprd positive afferents innervating the skin in diabetic conditions and responding with oscillatory calcium waves when activated with their known agonist β-alanine at the periphery could represent regenerating fibers. Recently Kuner and colleagues have elegantly revealed the emergence of a form of chronic neuropathic pain that is driven by structural plasticity, abnormal terminal connectivity, and malfunction of nociceptors during reinnervation^105^. Similar mechanisms could be at play in PDN. To answer this intriguing question mouse genetic approaches could be applied in future studies.

Regardless, our studies introduce the novel suggestion that Mrgprd is an important upstream mediator driving DRG neuronal hyperexcitability underlying mechanical allodynia in the HFD model. Hence, we propose Mrgprd and Mrgprd expressing neurons as promising accessible targets for developing effective therapeutics currently not available to treat neuropathic pain in diabetic neuropathy. Moreover, understanding which DRG neurons cell type is mediating mechanical allodynia in PDN is of fundamental importance to our basic understanding of somatosensation and may provide an important way forward for identifying cell-type-specific therapeutics to optimize neuropathic pain treatment and nerve regeneration in PDN. Thus, these results have the potential for transforming the way neuropathic pain in diabetic neuropathy is treated, replacing the largely ineffective approaches that are currently available for patients afflicted with PDN^13^. Furthermore, as demonstrated here, an unbiased approach combining single-cell transcriptomics and in vivo calcium imaging is a useful approach for revealing interesting targets that could be translated to produce more effective, disease-modifying therapies for patients suffering from PDN.

## MATERIALS AND METHODS

### Animals

All methods involving animals were approved by the Institutional Animal Care and Use Committee at Northwestern University. Animals were housed with food and water *ad libitum* on a 12-hour light cycle. We utilized the following mouse lines: C57/Bl6J (wild-type), Na_v_1.8-Cre;GCaMP6s, Mrgprd-eGFP reporter mice (MrgprdΔ EGFPf)^48^. Mrgprd-CreERT2 mice^68^ crossed with hM4Di mice^66^. The resulting Mrgprd-CreERT2;hM4Di mice were treated with tamoxifen 0.5 mg (Sigma-Aldrich, St. Louis, MO, T5648) given dissolved in corn oil via i.p. injection once per day starting postnatally day ten (P10) through postnatally day seventeen (P17). Animals were given at least one week to drive recombination and reporter gene expression.

### High-fat-diet

Mice were fed a diet with a high fat content (42% fat) (Envigo TD88137), High-fat-diet (HFD) for ten weeks as previously described^6,22,49,106^. Control mice were fed a regular diet (RD) containing 11% fat. After ten weeks on RD or HFD, a glucose tolerance test was performed as described^6,22^. Briefly, after fasting for 12 hours, mice were injected with a 45% D-glucose solution (2 mg glucose/g body weight). Animals were weighed on an electronic scale, and after fasting the animals for 12 hours, fasting blood glucose was measured using TrueTrack^®^ meter and TrueTrack^®^ glucose test strips. The mice were then injected with a 45% D-glucose solution (2 mg glucose/g body weight) and blood glucose was measured at 30, 60, and 120 minutes after injection (RD-Het n = 12 animals; RD-Homo n = 6 animals; HFD-Het n = 15 animals; HFD-Homo n = 15 animals). To compare “diabetic” versus “non-diabetic” HFD mice, we set the cutoff for diabetes (>178.41 mg/dL) at 2 SDs above the mean for glucose at 2 hours after glucose challenge, as determined from among wild-type littermate RD mice ^22^. **Statistics**: Blood glucose was analyzed using One-way ANOVA followed by Tukey’s test.

### Isolation of DRG neurons and single-cell RNA sequencing using 10X Genomics platform

Lumbar DRG neurons from adult mice fed on a regular diet (n = 5 animals) and a high-fat-diet (n = 5 animals) were isolated and dissociated following protocol established by Zeisel et al., 2018. A high viability single-cell suspension was prepared and using the 10X genomics chromium single cell kit v2, about 6000-8000 cells were recovered. Two rounds of the experiment (total n = 10 animals per group) were performed, and downstream cDNA synthesis, library preparation, and sequencing were performed according to the manufacturer’s instruction. Illumina runs were demultiplexed and aligned using the 10x genomics cell ranger pipeline. Dimensionality reduction, clustering and differential expression of genes were done in Seurat.

### RNAscope^®^ *in situ* hybridization

RNAscope^®^ in situ hybridization multiplex V2 was performed according to the manufacturer’s instructions (Advanced Cell Diagnostics, ACD). DRGs were isolated in an RNase free manner and the samples were then fixed in RNase free 4% PFA for 24 hours. The samples were transferred to 30% sucrose for 24 hours and embedded in OCT. 12μm sections were placed on SuperFrost Plus charged slides and stored at -20°C until ready to use. The slides were briefly washed in 1X PBS and followed by a 10-minute hydrogen peroxidase treatment at RT. The slides were then placed in a beaker containing 1X target retrieval solution which heated to around 99-102°C for about three minutes. The slides were then cooled in DEPC-treated water and transferred to 100% ethanol for three minutes. The slides were around to completely air dry and hydrophobic barriers were drawn around the sections. The air-dried slides were placed on the HybEZ™ slide rack and about 5 drops of RNAscope^®^ Protease III was added. The slide rack was placed onto a pre-warmed humidity control tray and into the HybEZ™ oven at 40°C for 30 minutes. Probes Scn10a (C1, catalog 426011), Nefh (C2, catalog 443671), Mrgprd (C3, catalog 45692), Lpar3 (C1, catalog 43259), Trpv1 (C2, catalog 313331), Human MRGPRD (C1, catalog 524871), and Human TRPV1 (C2, catalog 415381) were used at the recommended concentration (C1:C2; 50:1). Probes were incubated for 2 hours at 40°C and the slides were then stored in 5X saline sodium citrate solution. On day two, AMP1, AMP2 and AMP3 were added sequentially with a 30-, 30-, and a 15-minute incubation period, respectively. Depending on the probe used, the appropriate HRP signals were developed. Briefly, 4-6 drops of HRP-C1, or HRP-C2 were added and incubated for 15 minutes at 40°C. This was followed by the addition of 1:100 dilution of TSA^®^ Plus fluorescein. The fluorophores were incubated at 40°C for 30 minutes and this was followed by addition of the HRP blocker. Washes were performed using 1X wash buffer as recommended. The slides were then mounted using Vectashield mounting media containing DAPI. Tissue sections were analyzed by imaging the whole DRG using confocal microscopy.

### *In vivo* calcium imaging

Animals were fed an RD or HFD diet for 10 weeks, then anesthetized by isoflurane and laminectomized, exposing the L4 DRG as described^58^. The experimental set-up and imaging were done as previously reported^58^. Briefly, the mouse was positioned under the microscope by clamping the spinal column at L2 and L6; body temperature and isoflurane were constantly maintained and monitored throughout the imaging period. Silicone elastomer (World Precision Instruments) was used to cover the exposed DRG and surrounding tissue to avoid drying^58^. A Coherent Chameleon-Ultra2 Ti:Sapphire laser was tuned to 920 nm and GCaMP6s signal was collected by using a bandpass filter for the green channel (490 nm to 560 nm). Image acquisition was controlled using PrairieView software version 5.3. Images of the L4 DRG were acquired at 0.7 Hz, with a dwell time of 4 μs/pixel (pixel size 1.92 × 1.92 μm^2^), and a 10x air lens (Olympus UPLFLN U Plan Fluorite, 0.3 NA, 10 mm working distance). The scanned sample region was 981.36 × 981.36 μm^2^. Anesthesia was maintained using isoflurane (1.5-2%) during imaging.

### Intradermal β-alanine and capsaicin administration

5μL of 100mM β-alanine or 10μM capsaicin was injected intradermally to the paw pad in hind paw of anesthetized mice in the set up described in the previous section. The needle was inserted into the paw for 10 seconds and any responses to the needle were not included in the analysis. After 10 seconds, β-alanine or capsaicin was released and the neurons that responded to β-alanine or capsaicin reported as the percentage of neurons that responded. Appropriate controls like saline and ethanol were used to rule out non-specific responses as a result of needle injections.

### Analysis of *in vivo* calcium imaging

Time series files were exported and further processed in Fiji (NIH). Any movement in the time series (on account of breathing) were adjusted using the template-matching plugin. Brightness and contrast were adjusted and cells (region of interest, ROI) were identified by looking for an increase in fluorescence during the stimulus application period, as previously reported^58^. The identified cells were then carefully marked and a custom macro (where changes in [Ca^2+^]_i_ were quantified by calculating the change in fluorescence for each ROI in each frame t of a time series using the formula: ΔF/F_0_ = (F_t_ – F_0_)/F_0_, where F_o_ = the average intensity during the baseline period prior to the application of the stimulus) was run followed by a Multi Measure Plugin to obtain the mean gray value of each ROI. Once the values were obtained, the ROIs that had a ΔF/F_0_ reading greater than 1 were included as a responding ROI/neuron (RD n = 7 animals, HFD n = 9 animals, average number of neurons imaged per DRG in RD = 197.5 ± 48.67 and HFD = 242.5 ± 69.16). To determine the percentage of responders, the total number of neurons imaged for each DRG was estimated by counting the number of neurons within a region of average density and extrapolating to the total DRG area^58^. The area under the curve (AUC) was calculated for each responding ROI in GraphPad Prism 8.3, and the mean total peak area was calculated for each animal. The maximum amplitude was calculated by taking the maximum ΔF/F_0_ value for each responding ROI and the mean was calculated for each animal. **Statistics:** Percentage responders, AUC and max amplitude were compared using a two-tailed unpaired t-test. Paired t test was performed to compare saline and β-alanine as well as ethanol and capsaicin responses in both experimental groups. Data reported as mean ± SEM.

### Detection of cutaneous innervation

Animals were fed a RD or HFD for ten weeks. The hind paws were harvested and fixed in Zamboni fixative for 24 hours, and then the overlying footpad skin was dissected, submerged in 30% sucrose solution for 24 hours, and embedded in Optimal Cutting Temperature Compound (OCT, Tissue-Tek). 30μm sections were cut on a cryostat and counterstained by mounting solution with DAPI (Hardset, Vectashield). **Confocal analysis**: Two to three separate sections from each animal were analyzed and three separate composite Z-stack images of skin from the hind paw were imaged using Olympus FV10i and the images were processed using Fiji. The epidermal-dermal junction was outlined by a blinded observer who also noted its length. Two blinded reviewers counted the nerves crossing this line using the Cell Counter plugin. Mean values of the counts from blinded reviewers were divided by the epidermal-dermal junction length to report IENF density. **Statistical Analysis:** Data were compared using unpaired t test. In all experiments (RD n = 4 animals; HFD n = 3 animals), values are expressed as median ± SD.

### Intraperitoneal injection with clozapine-n-oxide

CNO (10 mg/kg) (Sigma Aldrich) in 200 microliters of saline were injected with a 25 gauge to mice fed either a RD or HFD. Mice were tested for pain 1 hour and 4 hours after injection with either CNO (RD-CNO n = 6 mice; HFD-CNO n = 6 mice) or vehicle (RD-saline n = 7 mice; HFD-saline n = 6 mice).

### von Frey

von Frey behavioral studies were performed as described. Briefly, mice were placed on a metal mesh floor and covered with a transparent plastic dome where they rested quietly after an initial few minute of exploration. Animals were habituated to this apparatus for 30 minutes a day, two days before behavioral testing. Following acclimation, each filament was applied to six spots spaced across the glabrous side of the hind paw. Mechanical stimuli were applied with seven filaments, each differing in the bending force delivered (10, 20, 40, 60, 80, 100, and 120 mN), but each fitting a flat tip and a fixed diameter of 0.2 mm. The force equivalence is 100 mN=10.197 grams. The filaments were tested in order of ascending force, with each filament delivered for 1 second in sequence from the 1st to the 6th spot, alternately from one paw to the other. The inter-stimulus interval was 10-15 seconds^22^. The von Frey withdrawal threshold was defined as the force that evoked a minimum detectable withdrawal on 50% of the tests at the same force. Experimental procedures were designed to maximize robustness and minimize bias. Specifically, von Frey experiments were conducted using random experimental group assignments (diet and treatment). Investigators that performed von Frey tests and endpoint analysis were blinded to the experimental conditions. We have experience with randomized allocation and blinded analysis using this mouse model with sequenced numbering of mice at weaning^22^. **Statistical analysis**: The incidence of foot withdrawal was expressed as a percentage of six applications of each filament as a function of force. A Hill equation was fitted to the function, relating the percentage of indentations eliciting a withdrawal to the force of indentation. From this equation, the threshold force was defined as the force corresponding to a 50% withdrawal rate^22^. Data were compared by One-way ANOVA followed by Kruskal-Wallis test (RD-Het n = 10 animals, RD-Homo n = 8 animals, HFD-Het n = 17 animals, HFD-Homo n = 16 animals) and reported as mean ± SD.

### Dynamic brush assay

Dynamic mechanical hypersensitivity was measured as previously described^61^. Briefly, the animals were placed on a 7.25mm spaced grid and covered with a transparent plastic dome where they rested quietly after an initial few minutes of exploration. Animals were habituated to this apparatus for 45 minutes a day, two days before behavioral testing. The hindpaw of the animals were lightly stroked in the heel to toe direction with a paintbrush (4310L). Both the right and the left hindpaw were brushed alternatively and scored with one minute of resting time between the brush strokes. We used a graded scoring system where the animal received a 0 score if the animal did not respond to the brush stimulus. Animals received a score of 1 if the animal lifted the paw of the mesh and placed it back on the mesh immediately. Animals received a score of 2 if there was sustained lifting (> 2 sec) or if there was flinching or licking of the stimulated paw. **Statistical analysis**: Stroking was repeated for a total of 10 times for each animal and the scores were averaged. Data were compared by One-way ANOVA followed by Kruskal-Wallis test (RD-Het n = 23 animals, RD-Homo n = 13 animals, HFD-Het n = 24 animals, HFD-Homo n = 22 animals) and reported as mean ± SD.

---

**Supplemental Figure 1:**
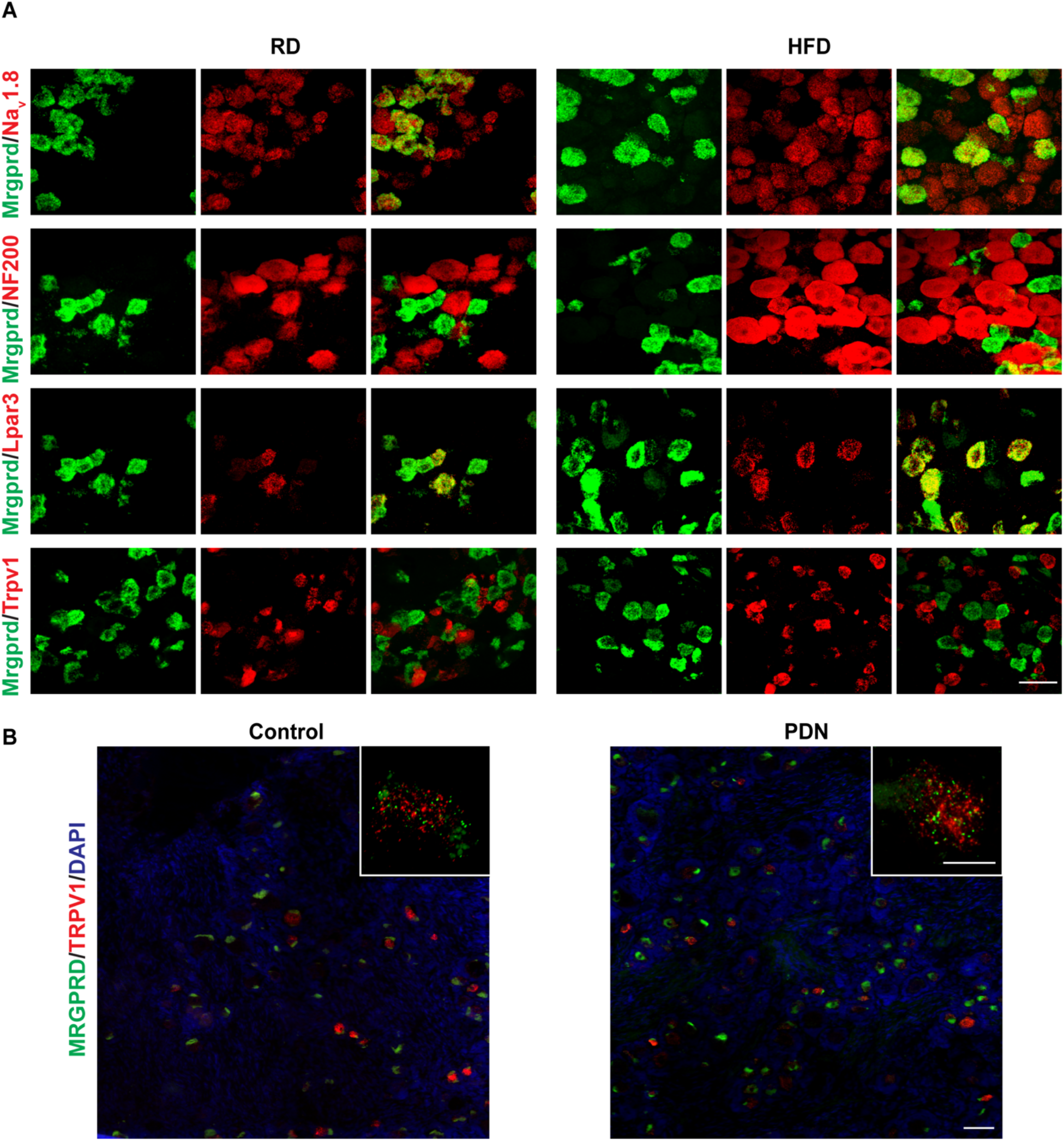
In-situ hybridization validating the expression of Mrgprd. **(A)** Na_v_1.8, Nefh, Lpar3, Trpv1 (red) with Mrgprd (green) from mice fed a RD or HFD. Scale bar represents 50μm. **(B)** MRGPRD (green), TRPV1 (red) and DAPI expression in human DRG samples from control and patients with PDN. Scale bar represents 200 μm. Inset shows a zoomed image of a single DRG neuron. Scale bar represents 25μm. RD n =3 mice and HFD n=3 mice; 3 sections were imaged for each group.

**Supplemental figure 2:**
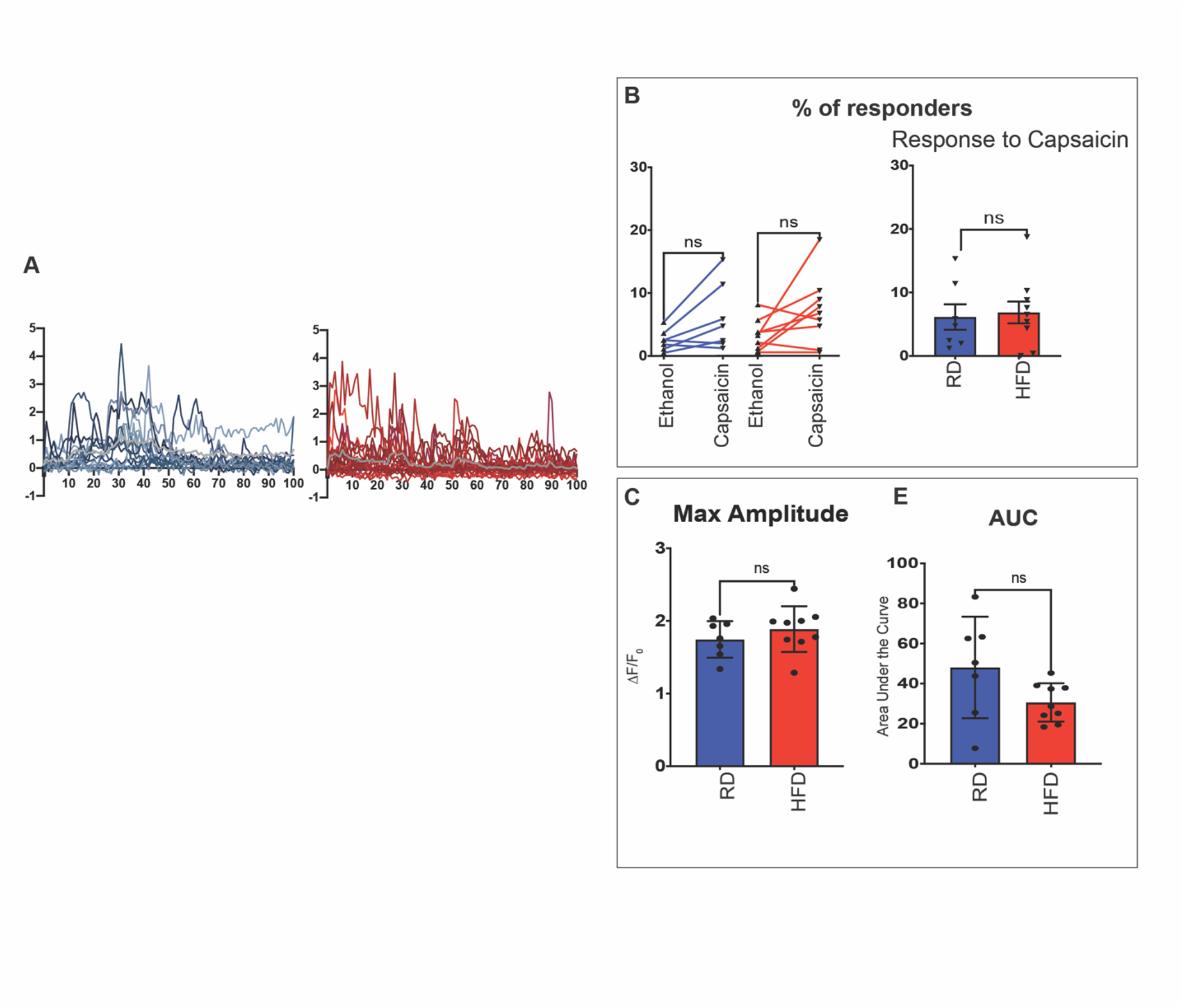
In vivo calcium response to capsaicin. **(A)** All responses to capsaicin from representative RD and HFD mice. The gray line indicates the average change in fluorescence intensity over time. **(B)** Paired analysis showing the percentage of responses to ethanol and capsaicin treatment in RD (blue) and HFD (red) and percentage of neurons responding to capsaicin, each black triangle represents one animal. **(C)** Graphs showing maximum amplitude and (**D**) area under the curve (AUC).

**Supplemental figure 3:**
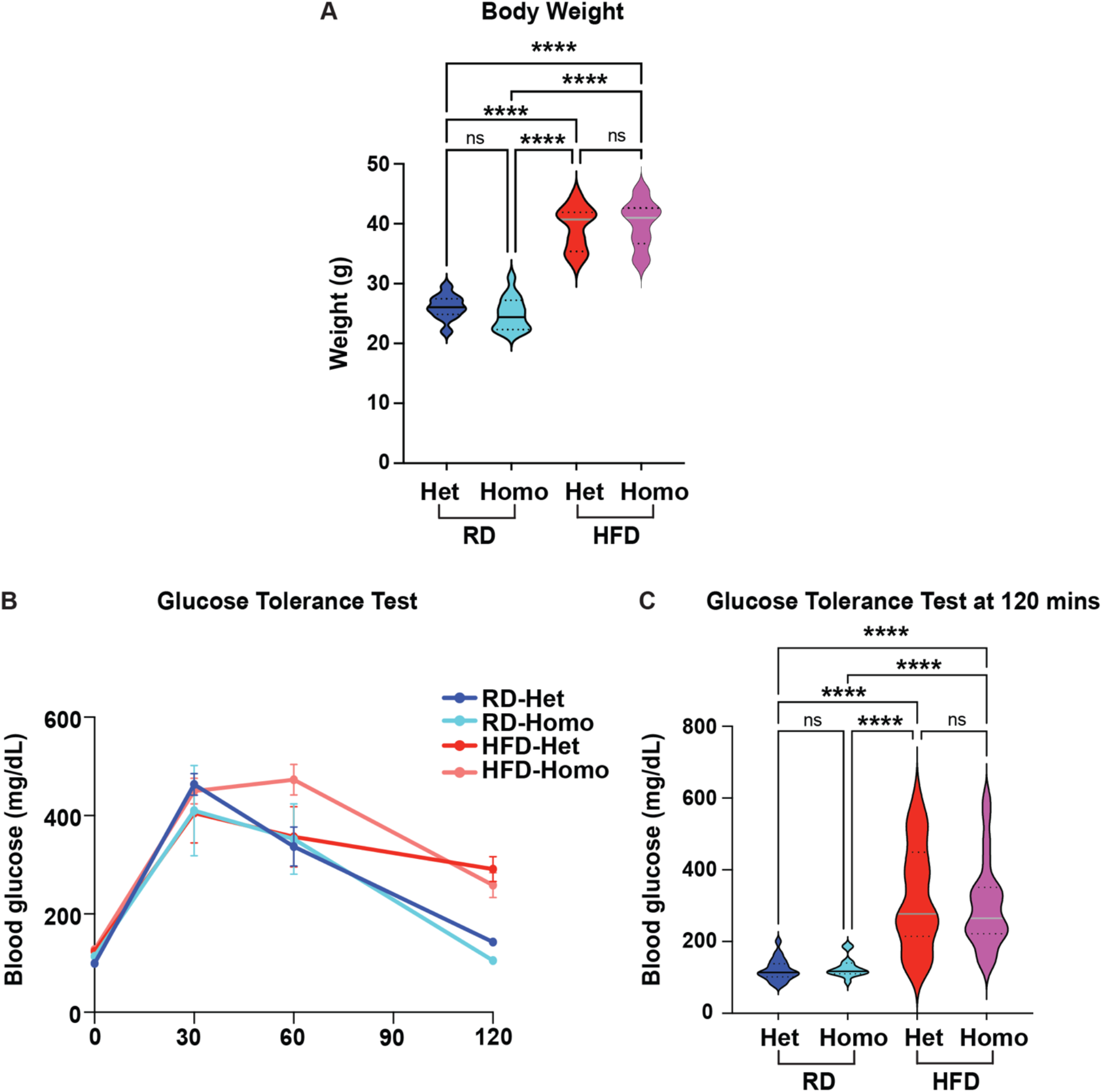
Deletion of Mrgprd does not alter the metabolic profile. **(A)** Body weight of Mrgprd heterozygous (+/-) and homozygous (-/-)mice fed either RD or HFD at ten weeks. **(B)** Blood glucose levels of the same mice at fasting, 30-, 60- and 120-minutes post glucose injection. **(C)** Area under the curve (AUC) for measurements in (B). RD-Het n = 10 mice; RD-Homo n = 8 mice; HFD-Het n = 17 mice; HFD-Homo n = 16 mice). ****p < 0.0001.

**Supplemental figure 4:**
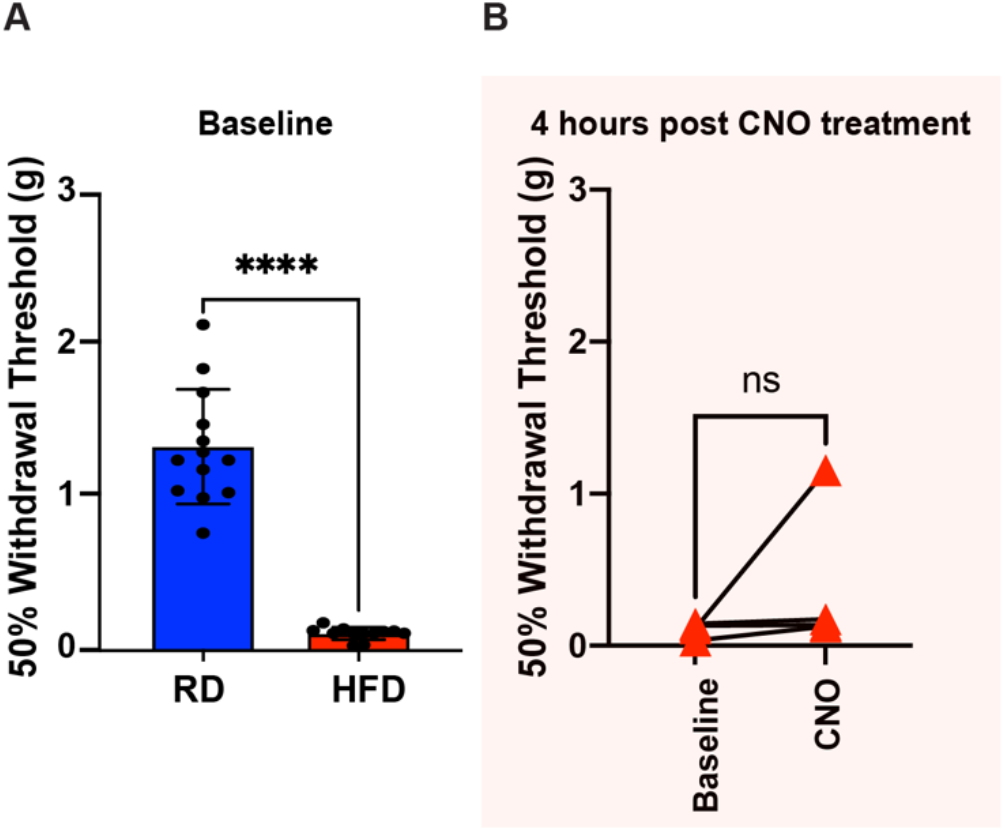
von Frey testing of RD and HFD Mrgprd-CreERT2;hM4D mice at **(A)** baseline and **(B)** four hours post CNO treatment. \*\*\*\**p* < 0.0001.

